# Analysis of the contribution of intrinsic disorder in shaping potyvirus genetic diversity

**DOI:** 10.1101/2022.05.13.491648

**Authors:** Guillaume Lafforgue, Thierry Michon, Justine Charon

**Affiliations:** INRAE; University of Sydney

## Abstract

Intrinsically disordered regions (IDRs) are abundant in the proteome of RNA viruses. The multifunctional properties of these regions are widely documented and their structural flexibility is associated with low constraint in the amino acid positions. Therefore, from an evolutionary stand point, these regions could have a greater mutational permissiveness than highly structured regions (ORs for Ordered Regions). They could thus provide a potential adaptive reservoir. To address this hypothesis, we compared the mutational robustness of IDRs and ORs in the genome of potyviruses, a major genus of plant viruses. For this purpose, a simulation model (DOI: 10.5281/zenodo.6396239) was built and used to distinguish a possible selection phenomenon in the biological data sets from randomly generated mutations. We analyzed several short-term experimental evolution datasets. An analysis was also performed on the natural diversity of three different species of potyviruses reflecting the long-term evolution. We observed that the mutational robustness of IDRs is significantly higher than that of ORs. Moreover, the substitutions in the ORs are very constrained by the conservation of the physico-chemical properties of the amino acids. This feature is not found in the IDRs where the substitutions tend to be more random. This reflects the weak structural constraints in these regions, in which an amino acid polymorphism is naturally conserved in the course of evolution, potyvirus IDRs and ODRs follow different evolutive paths with respect to their mutational robustness. These results force to consider the hypothesis that during selection, adaptive solutions could emerge from the amino acid polymorphism carried by IDRs.

## Introduction

### Protein intrinsic disorder

Proteins possess intrinsically disordered regions (IDRs), i.e. regions lacking a unique three dimensional structure and yet capable of exerting important biological functions [1,2], which challenges the so-called “structure-function relationship” dogma. Although it is today quite admitted that intrinsically disordered hub proteins are key players in the cellular interactome, the involvement of intrinsically disorder (ID) in evolution is still under debate. An earlier study was aimed at comparing the structural features of single-domain small-proteins from hypothermophylic bacteria, archaea, mesophilic eukaryota and prokaryota, and RNA or DNA viruses, whose crystal structures were available [3]. It was concluded from this analysis that viral proteins and more particularly RNA virus proteins, display (i) higher stability upon simulations of mutation accumulation and (ii) lower inter-residues contact densities. This latter feature is a typical signature of intrinsic disorder. It has thus been proposed that the large intrinsic disorder content in viral proteins could contribute to efficiently buffer mutation effects [3,4]. This was experimentally shown in the case of the intrinsically disordered protein VPg from potyviruses [5]. This is strongly contrasting with non-additive/epistatic stability loss profile expected from ordered proteins as previously reported for a bacterial β-lactamase [6]. It is hence conceivable that low structural requirements in IDRs could lead to some mutational robustness, and in turn, to an easier way for exploring the mutational space, without dramatic impairment of the protein biological functions. For instance, this idea sounds especially relevant regarding RNA virus adaptation to the host. This could contribute to a rapid adaptation to environmental stresses, without excessive loss of fitness. There is no doubt that this question is of very general interest. Consequently, the high evolutionary potential of RNA viruses, and the high ID content in their proteins, set the basis for assessing the contribution of ID to the shaping of virus genetic diversity in a context of host adaptation. Plant-phytovirus pathosystems provide useful experimental models for studying these aspects [7].

*A*n *in silico* analysis unveiled a high ID content in the *Potyvirus* proteome both at inter- and intra-species scales [8]. This feature has been conserved during *Potyvirus* evolution, suggesting a functional advantage of ID. When comparing the evolutionary constraint (ratio of non-synonymous to synonymous substitution rates, dN/dS) between ordered and disordered regions within the proteome of different potyvirus species, IDRs display significantly higher dN/dS values than ordered regions (ORs), a finding that indicates a tendency of intrinsically disordered domains to evolve faster than more structured regions during potyvirus evolution [8]. Using the pathosystem PVY/pepper, we previously obtained the first *in vivo* experimental data supporting the hypothesis that IDRs could influence virus adaptability to the host [9], possibly by enabling a faster exploration of the mutational space, thereby allowing the virus to bypass the plant resistance. Indeed, a correlation was observed between the adaptive potential of the virus and the disorder content within the VPg viral protein.

To further assess this previously described role of IDRs on RNA virus adaptation, the present study aimed at analyzing whether the regions predicted as disordered in viral proteomes are more likely to evolve and accommodate amino acid substitutions (non-synonymous mutations) than more structured areas. Ordered and disordered region sequences from various potyvirus species were thus retrieved and compared for several adaptive parameters at two timescales of viral evolution, a short -term scale experimental evolution and a long-term evolution reflected by natural diversity.

The short-term scale data analyzed consisted in high-throughput sequencing (HTS) retrieved from three independent evolution experiments, i.e. PVY [10,11], and TEV [12]. HTS provides access to the complete genome sequences of all viral variants - including those that are in a minority - that make up a population [13]. By sequencing each individual genome from the viral population, it is thus possible to assess the genetic structures of evolving potyvirus populations and thus potentially address the processes that shape this genetic variability, and to a greater extend the evolvability of the viral population.

To evaluate the impact of disordered versus ordered region on potyvirus evolvability (i.e. mutational robustness) at a higher scale of evolution, genomic sequences from TuMV, TEV and PVY natural diversity were also retrieved.

To prevent bias in our analysis of the structural determinant on potyvirus evolution, a third dataset, corresponding to simulated data was also obtained. Briefly, potyvirus genomes were artificially mutated *in silico* according the viral replicase features, to mimic the genetic diversity obtained in the absence of selection and, among others, effects of protein structural determinants. Adaptive parameters of the resulting mutants were thus obtained and compared to those from the biological data.

## Material and methods

### Data sets

#### Disorder prediction

We scanned for disordered regions along potyvirus polyproteins using Predictor of Naturally Disordered Regions (PONDR-VLXT), an algorithm accessible through the Disprot server (http://disorder.compbio.iupui.edu/metapredictor.php) [14,15]. Parameters were set to “default” for ID score predictions.

#### Experimental dataset

The study of Cuevas et al 2015 (TEV 2015), [12] evolved the TEV on two different host, *Nicotiana tabacum* and *Capsicum annuum*, while the two others studies Kutnjak et al 2015, 2017 [10,11] (PVY 2015 and PVY 2017), used the PVY on *Solanum tuberosum*. Table S1 compiles all the resulting mutations of experimental datasets.

#### Natural diversity dataset

Datasets used contained 6 genomes of TEV isolates, 100 genomes of PVY isolates and 100 genomes of TuMV isolates. Corresponding genome accessions are listed in Table S2. These datasets will be referred to as TEV_ND_, PVY_ND_ and TuMV_ND_ in the study.

#### Simulation

The distribution of mutations in the virus sequence is the sum of the contribution of viral polymerase errors and of the subsequent selection according to structure-function relationships. In order to uncouple these two components, we built an algorithm to mimic the distribution of synonymous and non-synonymous mutations introduced by the low fidelity virus RNA polymerase during genome replication (DOI: 10.5281/zenodo.6396239). It was hypothesized that, mutations could be randomly introduced all along the genome during its replication. Consequently, if IDRs and ORs were equally susceptible to mutations, NS and S were expected to be homogenously distributed in each of the two regions before virus submission to the selection pressure. The simulation takes also into consideration the specificity of viral polymerase on transversion/transition mutations calculated from TEV experimental data [16].

We generated n variants from the original potyvirus sequence, with each variant bearing one SNP.

#### Adaptive components tested

The collections of sequences generated from experimental evolution, natural diversity and simulated experiments where then analyzed with respect to the number of S and NS mutations (DOI: 10.5281/zenodo.6396239) and for each viral protein, their location either in the ORs or IDRs. BLOSUM-based scores of each NS mutations were also used to determine the potential of IDRs and ORs to cope with amino acid substitutions (DOI: 10.5281/zenodo.6396239). Finally, the characteristic of naturally occurring substitutions were analyzed in term of maintenance versus disturbance of disorder. It was reported that ORs and IDRs possess distinct sequence biases. Promotor scores, ranging from 0 to 1 (1 being the highest promoter score for disorder) were adapted from a previously published classification [17] and associated to each amino acids (Table S3).

## Results

To evaluate the contribution of intrinsically disordered regions on potyvirus evolvability and adaptation, this study compared viral genomic populations retrieved from evolution experiments, natural diversity and *in silico* generated pool of variants. Several parameters were thus assessed and compared at both genomic and proteomic levels and consisted in (i) the location of the diversity (within intrinsically disordered versus ordered protein or regions), (ii) the nature of nucleotide mutations (synonymous versus non-synonymous) as well as (iii) the biochemical and disorder-promoting nature of corresponding amino acid substitutions.

### Theoretical minimum number of mutations required for an accurate estimation of S and NS distribution in the genome

Datasets generated from the three experimental evolutions represent between 115 and 317 mutations. We hypothesized that the number of mutations considered could be too low to lead to robust conclusions. Consequently, a first consideration was to estimate the average number of mutations required to be significant. Four independent generations of 100, 300, 500, 750, 1000 and 1250 mutations were thus randomly introduced along the TEV genome sequence. Assuming that such random mutagenesis should not be impacted by any structural or protein determinants, the number of mutations (synonymous and non-synonymous) should be equally distributed along the genome, regardless of the corresponding proteome intrinsic disorder. Thus, the distribution of NS and S mutations among IDRs were determined (Figure 1). Above 600 mutations, an equal distribution of mutations among either ORs and IDRs was observed. Therefore, in order to ensure representative values for further analysis, the results of 4 independent simulations with 1000 mutations each, will be used.

**Figure 1.**
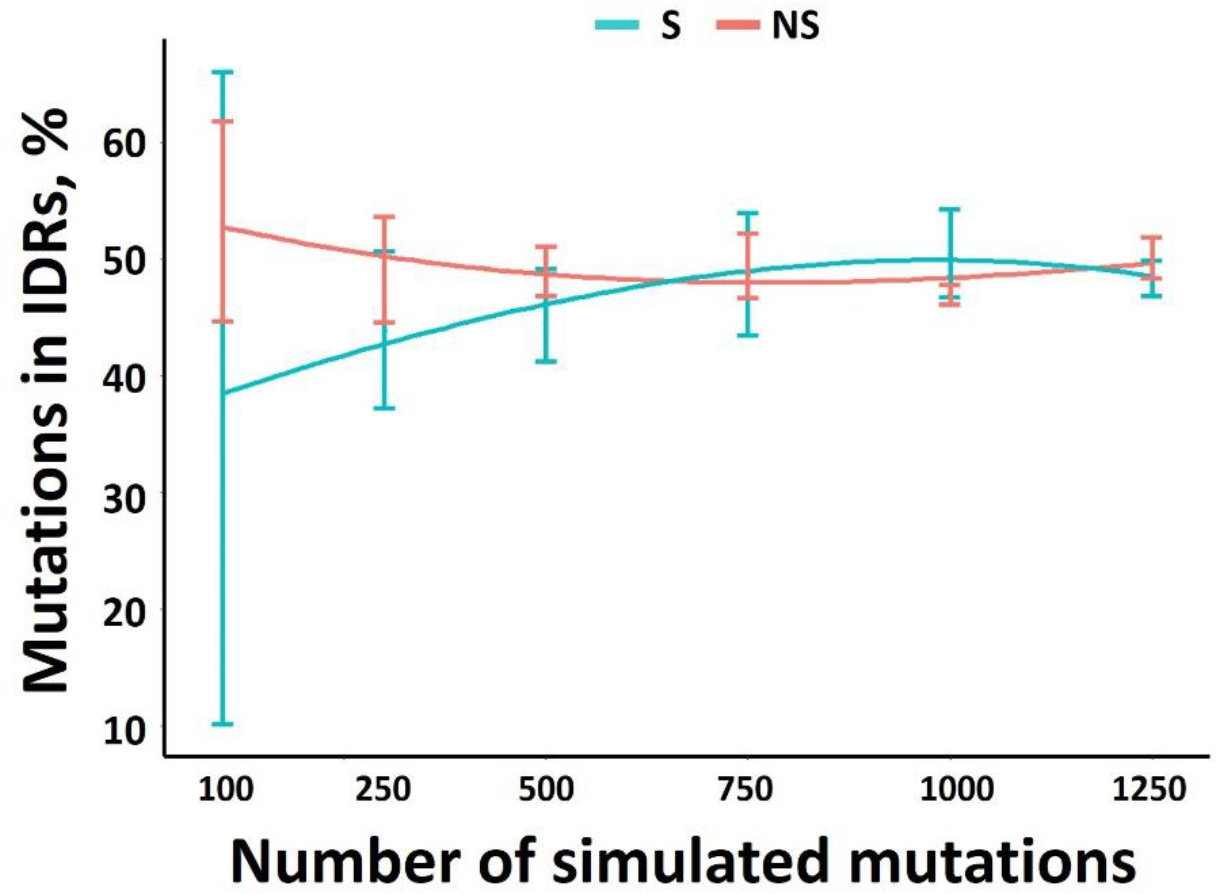
The % of S and NS mutations in IDRs versus the mutations number in the TEV genome. For a given number of mutations, 4 independent simulations were run.

This threshold of 1000 mutations, which is required for a robust analysis, was confirmed by monitoring the evolution of the R^2^ coefficient as a function of the mutation number. Whether for NS or S, below 750 mutations, the R^2^ greatly fluctuates (Figure S1).

This result confirms that the limited number of mutations available from the experimental data sets is likely to make our analysis less robust. To increase the size of the dataset and extend our observations to larger scales of viral evolution, the natural diversity of TEV_ND_, PVY_ND_ and TuMV_ND_ isolates was also analyzed by retrieving complete genomes available in Genbank. With 1296, 4646 and 7528 mutations identified in the corresponding TEV_ND_, PVY_ND_ and TuMV_ND_ datasets. These data should allow us to assess whether there is a significant difference in mutational robustness between IDRs and ORs.

### Correlation assessment between protein length and number of mutations

We first assessed the propensity of each potyvirus proteins to accumulate adaptive non-synonymous (NS) versus synonymous (S) mutations. The number of NS or S mutations observed in each protein coding sequence divided by the total protein length were thus calculated for each of the experimental evolution, natural diversity and simulated data sets.

At the short-term evolution scale, the longer the protein, the higher the number of S mutations, with a significant correlation between protein length and percentage of S mutations. By contrast NS mutation number were not correlated with the protein length, for the three experimental studies analyzed [10–12] (Table 1).

**Table 1.** Correlation coefficient (R^2^) between coding sequence length of the TEV proteins and the mutations (S or NS). Experimental evolution: TEV 2015 [12], PVY 2015 [10] and PVY 2017 [11]. TEV_ND_, PVY_ND_, TuMV_ND_: ND natural diversity. Simulations: four *in silico* replicates.

|  | S | NS |
| --- | --- | --- |
| TEV 2015 | 0,63 | 0,19 |
| TEV<sub>ND</sub> | 0,94 | 0,16 |
| Simulations | 1 | 0,98 |
| PVY 2015 | 0,93 | 0,12 |
| PVY 2017 | 0,78 | 0,01 |
| PVY<sub>ND</sub> | 0,96 | 0,35 |
| Simulations | 0,96 | 0,97 |
| TuMV<sub>ND</sub> | 0,95 | 0,09 |
| Simulations | 0,92 | 0,98 |

At the long-term evolution scale, the natural diversity confirmed the trend that the accumulation of NS poorly correlated with protein length. Non-synonymous adaptive mutations, which reflect the viral amino acid polymorphism, are thus not accumulated homogenously along the potyvirus proteome.

Regarding the simulated data, S and NS mutations are equally represented along potyvirus mutated genomes, independently of the protein length (R^2^≃0.94), thus validating our random model for a number of mutations above 1000. As expected, it is indicative of the correlation between the protein sequence length and the number of S or NS mutations obtained at random in the absence of any biological bias.

To be noticed, for all simulated data, the number of NS mutations is 3 times higher than the number of S mutations. By contrast, for the experimental data, the number of S and NS mutations is equivalent. Assuming that there is little or no selection pressure on S mutations, we can extrapolate the number of NS mutations before selection. So, in the TEV 2015 experiment [12], the total number of mutations before selection would be 300 mutations distributed in 75 S and 225 NS mutations, 278 mutations for the PVY 2015 experiment [10] and 1077 mutations for the PVY 2017 experiment [11]. We can see that for two datasets, before selection, we do not reach the minimum required of 1000 mutations previously defined to obtain a robust analysis.

The amount of S is rigorously proportional with the protein length irrespective of its function. This is not the case for NS, and some proteins contain more mutations then others. For instance, it appears that P1 significantly accumulates more NS than HC-Pro, P3, CI, NIb and CP (P < 0.02; Z test) (Figure 2 and Figure S2).

**Figure 2.**
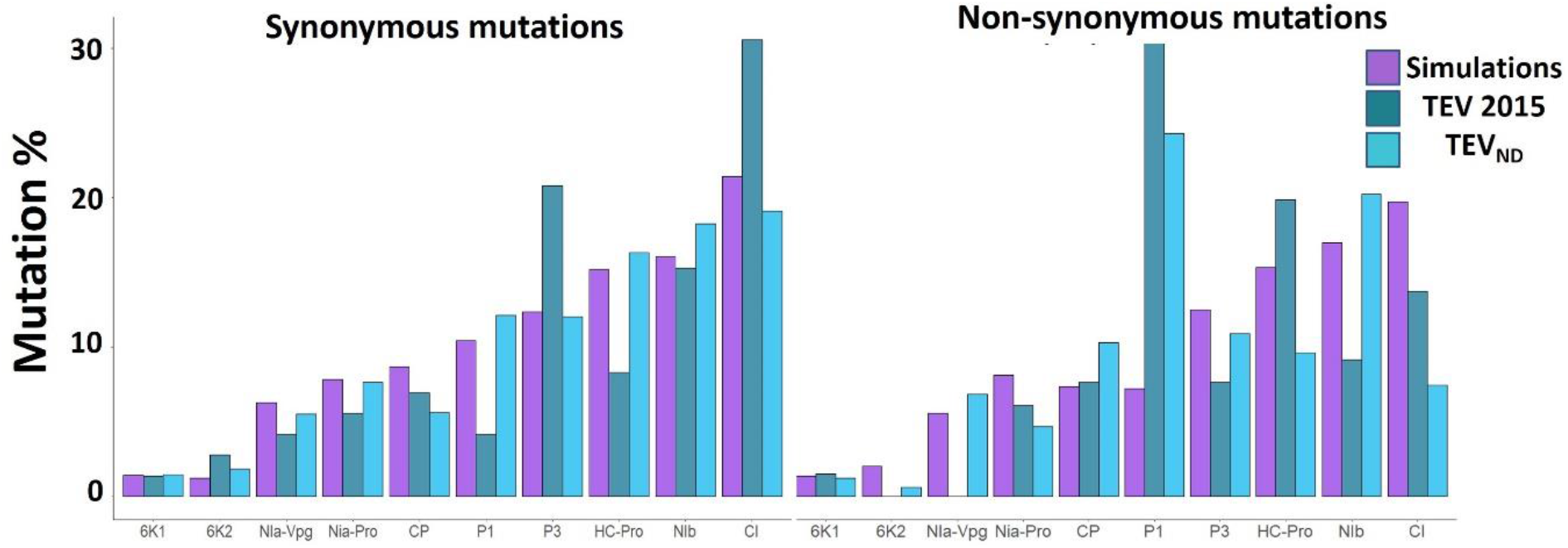
Mutation % in the TEV proteins from the experimental evolution [12], natural diversity and simulations. For PVY and TuMV see supplemental data. The proteins are sorted from the smallest to the largest, left to right: 6K1, 6K2, Nia-VPg, Nia-Pro, CP, P1, P3, Hc-Pro, Nib, CI.

### DISTRIBUTION OF MUTATIONS NS and S in IDRs and ORs

In the second part of the study, the analysis was no longer conducted on individual proteins, but on all IDRs and ORs distributed along the coding sequences in the viral genomes. In order to analyze the distribution of each type of mutation in the IDRs or ORs, we defined the ratio *R* for synonymous mutations as:

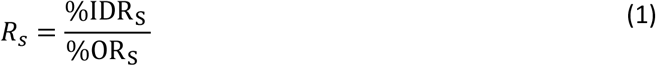

with %IDR_S_ and %OR_S_ defined as

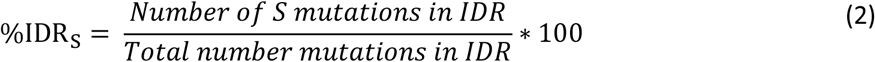

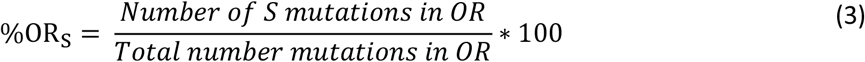

Equations 1-3 also apply for the calculation of *R*_NS_, the ratio R for non-synonymous mutations.

The ratio of synonymous mutations between IDRs and ORs deduced from the experimental data were close to 1 (Figure 3 and Figure S3). This ratio is comparable to that obtained by simulation which mimics random mutations and reflects the absence of impact of synonymous mutations at the protein level. By contrast, for NS mutations, a large and significant difference between the experimental and simulated data could be observed (p<0.02, χ2 for each of the four simulations, Table 2). Indeed, the ratio higher than 1 observed in the case of the experimental data indicates an over-representation of NS mutations within the IDRs compare to the ORs (Figure 3). In the case of PVY, this difference with simulated mutations was only verified for the PVY 2017 dataset [11] (Table 2).

**Table 2.** p values of Xhi^2^ test for percentage of NS mutations in IDRs between simulated and experimental data (TEV 2015, PVY 2015, PVY 2017) or natural diversity (TEV_ND_, PVY_ND_, TuMV_ND_). There was no experimental datasets available for TuMV. Significance, p<0.05.

| Simulations | TEV 2015 | TEV<sub>ND</sub> | PVY 2015 | PVY 2017 | PVY<sub>ND</sub> | TuMV<sub>ND</sub> |
| --- | --- | --- | --- | --- | --- | --- |
| A | 0,014 | 2.10-6 | 0,52 | 0,04 | 0,0005 | 3.10-8 |
| B | 0,026 | 1.10-5 | 0,46 | 0,03 | 0,0001 | 5.10-8 |
| C | 0,018 | 5.10-6 | 0,33 | 0,01 | 8.10-6 | 2.10-8 |
| D | 0,028 | 2.10-5 | 0,27 | 0,0049 | 1.10-7 | 6.10-9 |

**Figure 3.**
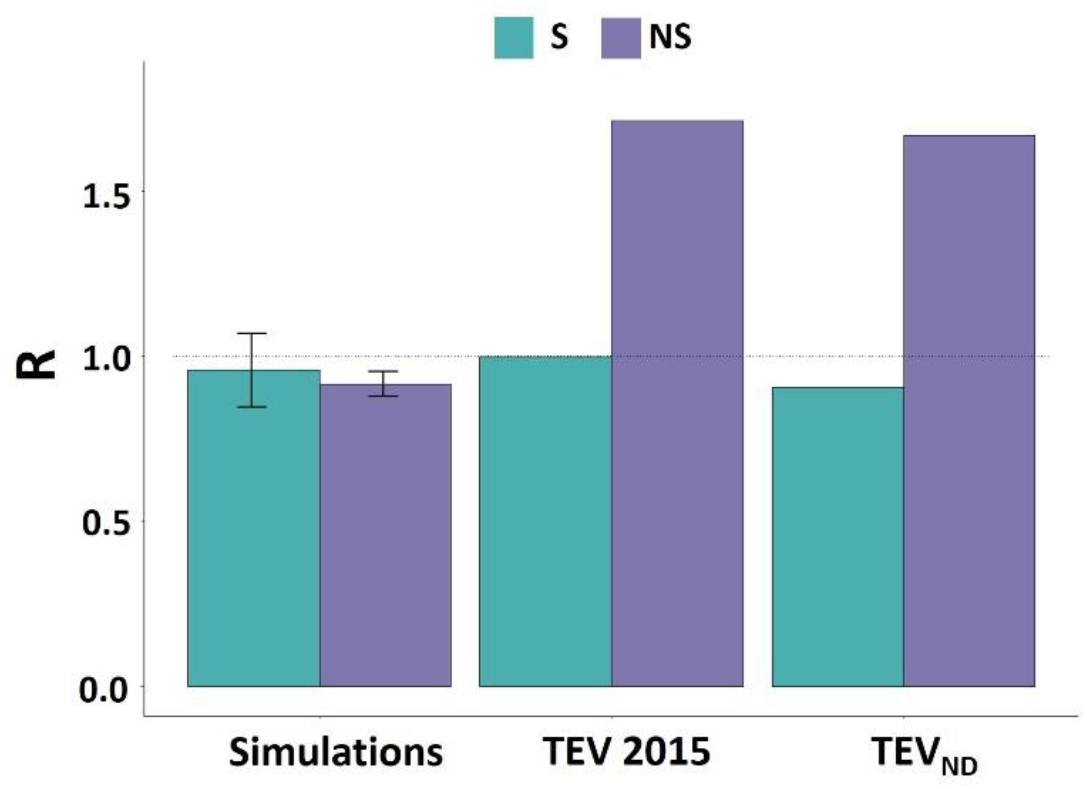
Ratio between the percentage of mutation (S or NS) present in IDRs and ORs for the TEV genome. For data sets from the two other studies [10,11], TuMV and PVY see supplemental data.

With respect to the analysis of natural diversity, no differences were observed between S mutations within IDRs and ORs, in agreement with simulated data. By contrast, a significant over-representation of NS was observed in the IDRs for all viruses (Figure 3 and Figure S3).

Altogether, those results indicate that IDRs are more prone to accumulate adaptive mutations than more structured regions, at both short (experimental evolution) and longer (natural diversity) evolutionary time-scales.

### Comparison of the physicochemical disturbance of amino acid substitutions in potyviral intrinsically disordered versus ordered regions

We also analyzed possible amino acid substitution biases between IDRs and ORs with respect to their physico-chemical properties. To compare the physico-chemical nature of the substitutive amino acids (NS mutations) in IDRs and ORs, we used the BLOSUM62 matrix [18]. This matrix uses the natural diversity between very conserved regions of evolutionary divergent protein sequences. The set of sequences is aligned to a reference sequence. For each position where a substitution occurs, the probability of occurrence of each of the 19 other amino acids is calculated, resulting in score values ranging between -4 and 11. The higher the score, the higher is the likelihood of substitution. It was observed that the highest replacement probabilities is correlated to amino acids with similar physico-chemical properties (charges, hydrophilicity-hydrophobicity, amino acid size) [19]. Upon amino acid substitutions, the more drastic physicochemical changes are (the lower the score value), the more destabilizing these changes are in terms of structure.

For each virus (TEV, PVY, and TuMV), we assessed whether the natural selection discriminated differently within IDRs and ORs for amino acid substitutions with respect to their impact on biophysical changes. For each type of region (IDRs or ORs), a comparative statistical analysis (Dunn test) was thus performed between the natural diversity, experimental and simulated data sets (Table 2 and Table S4). When amino acid substitutions occur in IDRs, their BLOSUM62 scores in the simulated data and those in the PVY and TuMV data sets belong to the same statistical group. In the case of TEV, the natural diversity data shows a slight difference with three of the four simulations. Importantly, its diversity was represented by a set of only 6 genomes while those of PVY and TuMV were illustrated by 100 genomes each. In contrast, for all three potyviruses, regarding the amino acid substitutions present in the ORs, both natural diversity and experimental evolution data have a significantly higher BLOSUM62 score than the simulated data. The high BLOSUM62 score observed as associated to ordered regions supports the idea that amino acids substitutions occurring in those regions are globally poorly destabilizing at the physicochemical and structural level. Reciprocally, with lower BLOSUM62 scores than the ones observed in ORs, IDRs would be more permissive to drastic physicochemical changes.

**Table 2.** Differences in physicochemical properties associated with amino acid substitutions were assessed using scores derived from the BLOSSOM62 substitution matrix. For each type of region (IDRs or ORs) groups (a,b and c) were determined by running a Dunn test (p value adjustment method: Bonferroni). For (A) PVY genome, (B) TEV genome and (C) TuMV genome.

| A | Groups |  |
| --- | --- | --- |
|  | IDRs | ORs |
| PVY 2015 | abc | ab |
| PVY 2017 | abc | a |
| PVY<sub>ND</sub> | ac | a |
| Sim A | bc | c |
| Sim B | bc | c |
| Sim C | b | bc |
| Sim D | bc | c |

| B | Groups |  |
| --- | --- | --- |
|  | IDRs | ORs |
| TEV 2015 | ab | ab |
| TEV<sub>ND</sub> | b | b |
| Sim A | a | a |
| Sim B | a | a |
| Sim C | a | a |
| Sim D | ab | a |

**Table 2.** Differences in physicochemical properties associated with amino acid substitutions were assessed using scores derived from the BLOSSOM62 substitution matrix.
| C | Groups |  |
| --- | --- | --- |
|  | IDRs | ORs |
| TuMV<sub>ND</sub> | a | a |
| Sim A | a | b |
| Sim B | a | b |
| Sim C | a | b |
| Sim D | a | b |

### Are NS mutations in IDRs driven toward the conservation of disorder promoting amino acids ?

We investigated whether the conservation of disorder during evolution could be a selection criterion using amino acid disorder promoting scores (see material and method section). Amino acid residues were grouped into order promoting, neutral or disorder promoting scores, ranging from 0 (amino acid most frequently present in ORs) to 1 (amino acid most frequently present in IDRs), were attributed to each of the 20 amino acids [17,20].

We first examined if, substitutions were preferentially targeting order or disorder promoting amino acids, and this, whether in IDRs or ORs. We did not observe any significant differences between biological data within either IDRs or ORs and simulated data (Table S5-A). We concluded that there is no natural tendency for evolution to target substitutions preferentially toward order or disorder promoting amino acids.

Then, we considered the possibility that non-synonymous mutations (NS) could preferably give order or disorder promoting amino acids (Table S5-B). We observed unbiased random substitution, and this, both in ORs or IDRs, in accordance with simulations. Finally, we aimed at assessing a possible tendency for substitution by amino acids that are more prone to promote order or disorder (Table S5-C). At each position where a NS mutation was observed, we calculated the difference in promoter score between the amino acid in the reference genome and the replacing amino acid in each of the genomes describing the diversity in the biological data. Again, we did not observe significant differences between naturally selected and simulated mutations. We did not detect any differences between biological and simulated data in the promoter score for the synonymous mutations, either in the IDRs or ORs. However, global disorder is generally conserved during evolution [21–23], and more specifically in RNA viruses [4,8]. Therefore, the analysis of local substitutions does not reflect this evolutionary trend that can be observed globally at the scale of a protein region. It turns out that the analysis of substitutions in terms of physico-chemical modulations sounds more relevant than the use of the order-disorder promoter scale.

## Discussion

### Mutational robustness differences between IDRs and ORs

Potyviruses constitute, together with begomoviruses, the two largest viral genera described to date among plant viruses [24,25]. Potyviruses are very damaging to field crops and embrace a very wide host range [26]. Most of them are generalists and as such, provide a rich model for studying viral adaptation. In this study, we tested the hypothesis that among these viruses, mutational robustness was greater in the disordered regions of their proteomes than in the ordered regions. We analyzed the distribution of mutations in the genomes of potyviruses belonging to three different species, PVY, TEV and TuMV, representative of the genus. The datasets used included viral genomes resulting from both short evolutionary scale (experimental evolution) and longer evolutionary scales, with the use of natural diversity. An analysis of the two datasets showed that IDRs and ORs are subject to different evolutionary mechanisms, with disordered regions evolving towards significantly more amino acid polymorphism than ordered regions. The selection pressure that applies to ordered regions thus tends towards a conservative evolution while that which applies to disordered regions rather supports a divergent evolution. It is quite easy to understand the evolutionary mechanism at work in the ordered regions. These protein regions have a strong structure-function relationship. At the molecular level, these regions are defined by geometries of constrained atomic interactions with few degrees of freedom and well-packed hydrophobic cores. These regions have significantly higher BLOSUM62 scores than would result from random substitutions. Substituted amino acids have physicochemical natures close to those of the original amino acids. Conversely, the lower topological requirement in disordered regions results in substitutions close to the random substitution pattern. Within ordered regions, mutations compensate each other to prevent instability according to an epistatic model. On the long term, such compensation leads to changes in sequence and function (protein evolvability) [6,27]. In these regions the selection pressure strongly operates to preserve function which results in amino acids conservation. In disordered regions, the notion of structural stability is less relevant and amino acids substitutions may have less functional impact [4]. This could constitute an alternative model for protein evolvability, presumably on a shorter evolutive timeline consistent with the rapid adaptation characteristic of viruses. From an evolutionary standpoint, the dogma of the structure-function relationship (conservation of function requiring conserved structures and therefore close substitutions) requires to be tempered in the case of IDRs.

### Does amino acid polymorphism in potyvirus proteome IDRs undergoes positive selection?

We examined the hypothesis that intrinsic disorder could be selected to generate a pool of mutations available for adaptive function. To obtain the diversity observed in IDRs, two successive processes, namely the generation of mutations and their selection, are involved. The first process can be favored by codon volatility. Codon volatility is defined as the proportion of a codon’s point mutation neighbors that code for different amino acids [28]. We investigated whether the nucleotide sequences encoding IDRs used codons of higher volatility than the sequences of ORs, thus favoring the generation of non-synonymous mutations. We did not observe greater codon volatility in the disordered regions than in the ORs that could explain the greater amino acid polymorphism observed in the disordered regions. Thus, with respect to the volatility criterion, we have no evidence to support that amino acid polymorphism in disordered regions undergoes positive selection, generating a potential adaptive pool for the virus. Although amino acid polymorphism in these regions may participate in potyvirus adaptation, the conservation of intrinsic disorder during evolution is the result primarily of the second process, a selection pressure dictated by the essential biochemical functions it performs to ensure virus replication in the host. In any case, the mutational permissiveness and diversity that arise from the selection of structure-function relationships within IDRs is likely to favor the adaptive potential of the virus. It cannot therefore be excluded that IDRs are also selected according to this last criterion, even if this hypothesis remains difficult to assess.

### Evolutive features of nucleotide sequences encoding IDRs

It should be expected that the synonymous codon usage pattern of viruses would be shaped by selecting specific codon subsets to match the most abundant host transfer RNAs (tRNAs). However, the codon usage of many viruses is very different from the optimal codons present in the host [29]. Interestingly, it was recently reported that codon usage in virus IDRs is less optimized for the host than in ORs [30]. In the case of NS mutations, this is in line with our observation that IDRs are more robust to mutations than OR, and thus evolve faster. This prevents fixation of codons optimized for the host (figure 5). The preservation of codon diversity in these regions may also provide a reservoir for a faster adaptation of the viruses to various hosts.

**Figure 5.**
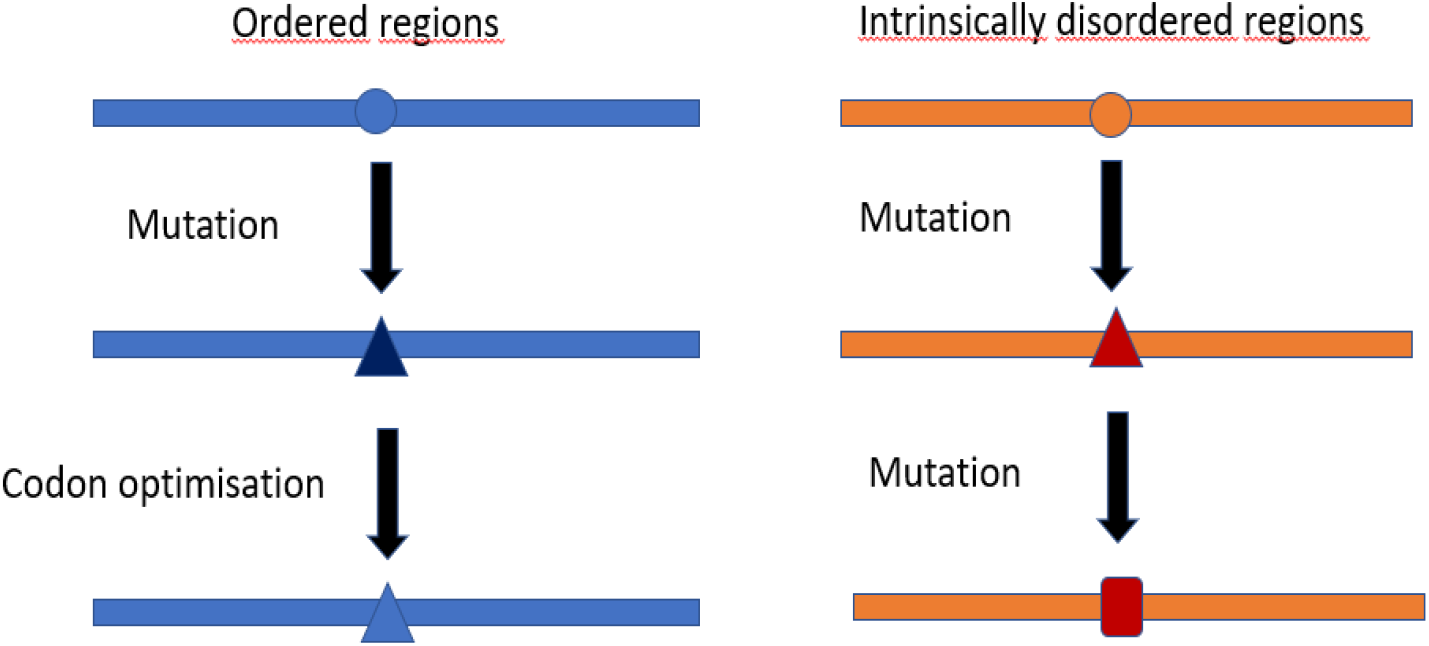
Mutational robustness of IDRs and codon optimization. The low rate of non-synonymous mutations in the ORs allows the optimization of the sequence towards the selection of abundant codons in the host. The high rate of non-synonymous mutations in IDRs prevents this optimization.

Because of the presence of less frequent codons in IDRs, the corresponding pools of loaded tRNAs in the host cell are lower than those of abundant codons. Consequently, translational dynamics is likely to be slowed down when the ribosome machinery enters a mRNA sequence encoding for disordered regions [31,32]. This may result in an instability of the translation product [33]. IDRs are generally taken over either co-translationally or post-translationally by chaperones. This handling does not favor the selection of optimized codons and contributes to the preservation of amino acid polymorphism in IDRs. There is an intricate interplay of molecular chaperones and protein disorder in the evolvability of protein networks [34].

Taken all together, the data obtained unambiguously show that potyvirus IDRs and ODRs follow very different evolutive paths with respect to their mutational robustness. These results force to consider the hypothesis that during selection, adaptive solutions could emerge from the amino acid polymorphism carried by IDRs.

## Supporting information

Lafforgue 2022- Supplemental data (Figures)

Lafforgue 2022 - Table S1

Lafforgue 2022 - Table S2

Lafforgue 2022 - Table S3

Lafforgue 2022 - Table S4

Lafforgue 2022 - Table S5

