## Supplementary material for "Analysis of the contribution of intrinsic disorder in shaping potyvirus genetic diversity": Lafforgue 2022- Supplemental data (Figures)

**Table S1.** Mutations in coding sequences detected during experimental evolution in PVY 2015 1. (A) and PVY 2017 2. (B) or in TEV 2015 3. C).

1. Kutnjak  et al. J Virol. 2015;89: 4760–4769. doi:10.1128/jvi.03685-14

2. Kutnjak et al. J Virol. 2017;91. doi:10.1128/jvi.00690-17

3. Cuevas et al. Mol Biol Evol. 2015;32: 1132–1147. doi:10.1093/molbev/msv028

**Table S2.** PVY, TuMV and TEV isolates used to generate the natural diversity dataset.

**Table S3.** Adapted promotor score
from Radivojac P et al. Biophys J. 2007;92: 1439–1456. doi:10.1529/biophysj.106.094045

**Table S4.** Differences in physicochemical properties associated with amino acid substitutions were assessed using scores derived from the BLOSOM62 substitution matrix

**Table S5.** Table S5, Dunn test (P value adjustment method : Bonferroni) of the order/disorder promoting score of each amino acid of the reference genome targeted by substitutions before substitution (A), of the amino acid resulting from substitution (B) and the differences in promoter score between the amino acid in the reference genome and the replacing amino acid in each of the genomes (C).

**Figure S1.** Variation of $R^2$, the coefficient referring to the correlation between percentage of mutations (S or NS) and protein length in the TEV genome, versus the mutations number. For a given number of mutations, 4 independent simulations were run.

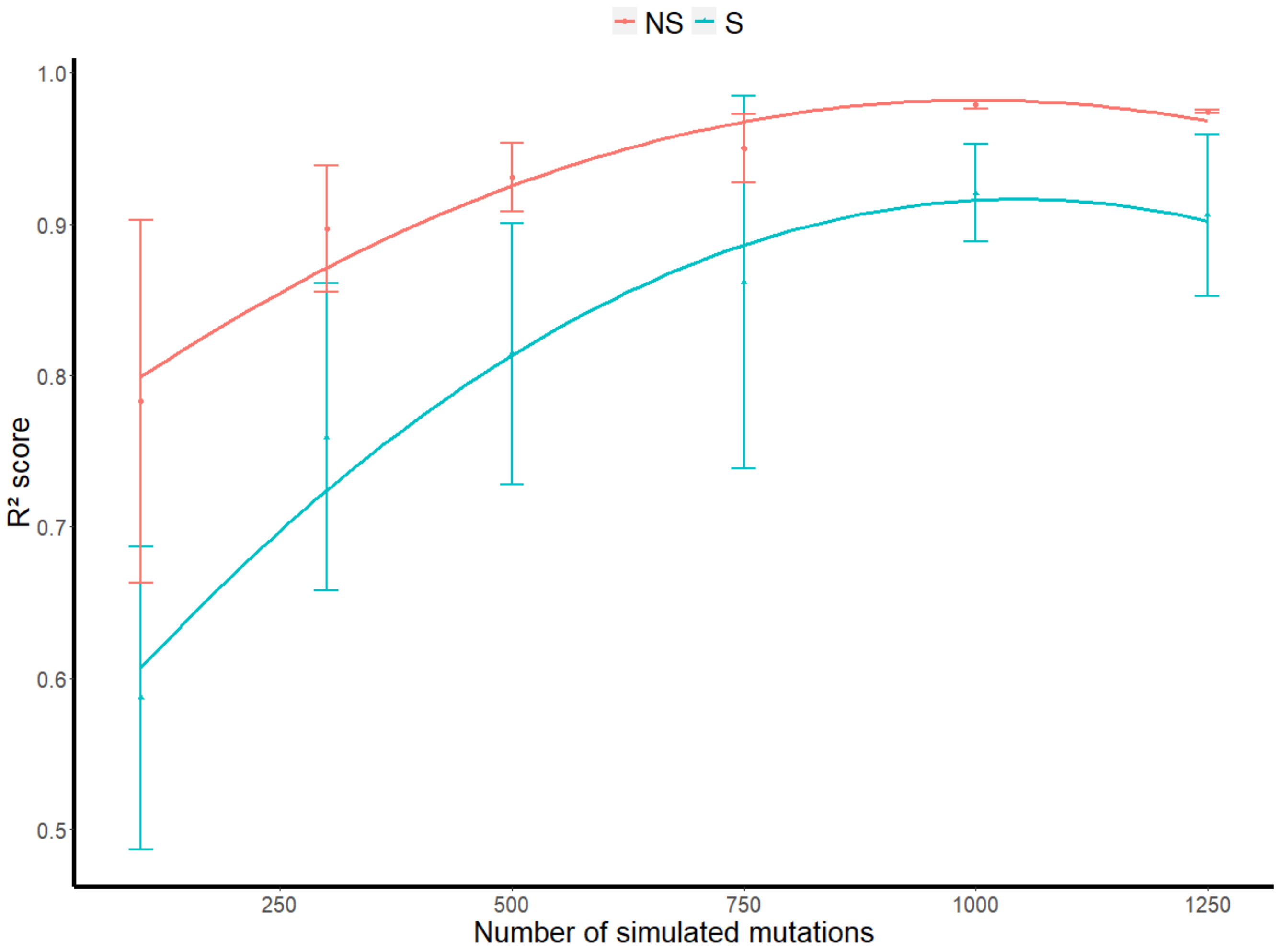

**Figure S2.** Mutation % in A. PVY proteins from experimental evolution (PVY 2015-2017), natural diversity ($PVY_{ND}$) and simulations. B. TuMV proteins natural diversity ($TuMV_{ND}$) and simulations. The proteins are sorted from the smallest to the largest, left to right: 6K1, 6K2, Nia-VPg, Nia-Pro, CP, P1, P3, HcPro, Nib, CI.

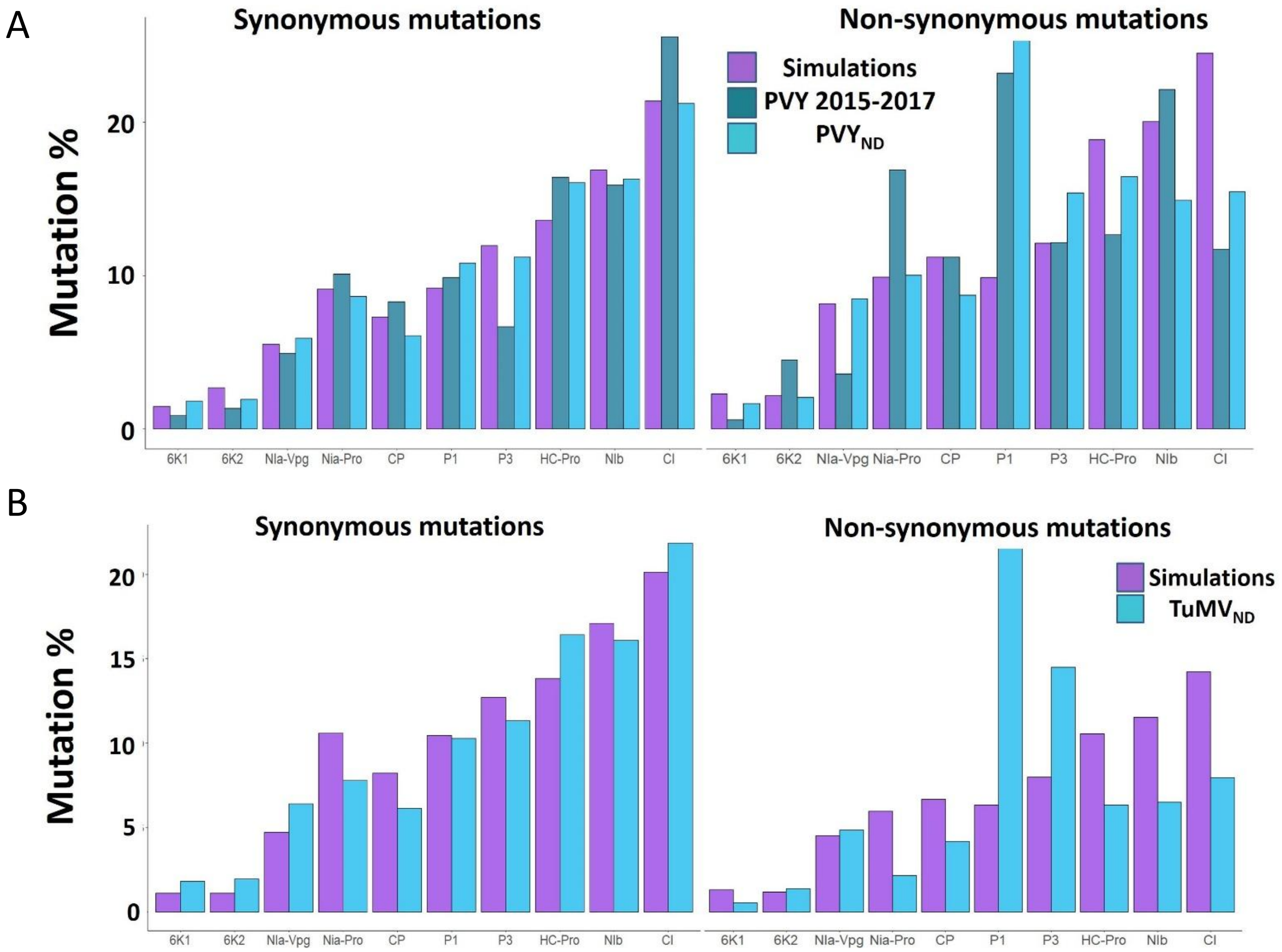

**Figure S3.** Ratio between the percentage of mutations (S or NS) present in IDRs and ORs for PVY (A), and TuMV (B). From left to right, data simulated (4 simulations), datasets from the experimental evolution (PVY 2015-2017) and natural biodiversity ($PVY_{ND}$ or $TuMV_{ND}$).

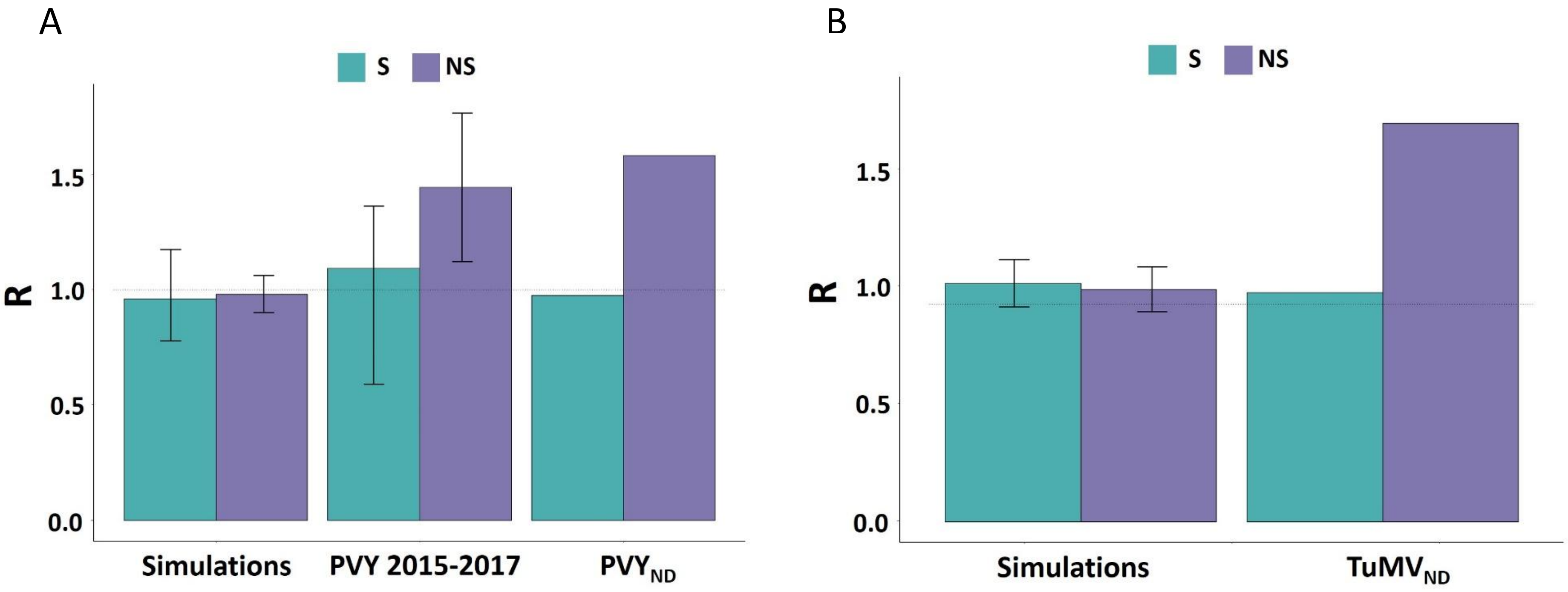
